# The social situation affects how we process feedback about our actions

**DOI:** 10.1101/428052

**Authors:** Artur Czeszumski, Benedikt V. Ehinger, Basil Wahn, Peter König

**Affiliations:** Institute of Cognitive Science, Universität Osnabrück, Osnabrück, Germany; University of British Columbia, Department of Psychology, Vancouver, BC, Canada; Institut für Neurophysiologie und Pathophysiologie, Universitätsklinikum Hamburg-Eppendorf, Hamburg, Germany

**Keywords:** Social Cognition, Joint Action, EEG, Feedback related negativity, Coopera-tion, Competition

## Abstract

Humans achieve their goals in joint action tasks either by cooperation or competition. In the present study, we investigated the neural processes underpinning error and monetary rewards processing in such cooperative and competitive situations. We used electroencephalography (EEG) and analyzed event-related potentials (ERPs) triggered by feedback in both social situations. 26 dyads performed a joint four-alternative forced choice (4AFC) visual task either cooperatively or competitively. At the end of each trial, participants received performance feedback about their individual and joint errors and accompanying monetary rewards. Furthermore, the outcome, i.e. resulting positive, negative or neutral rewards, was dependent on the pay-off matrix, defining the social situation either as cooperative or competitive. We used linear mixed effects models to analyze the feedback-related-negativity (FRN) and used the Threshold-free cluster enhancement (TFCE) method to explore activations of all electrodes and times. We found main effects of the outcome and social situation at mid-line frontal electrodes. The FRN was more negative for losses than wins in both social situations. However, the FRN amplitudes differed between social situations. Moreover, we compared monetary with neutral outcomes in both social situations. Our exploratory TFCE analysis revealed that processing of feedback differs between cooperative and competitive situations at right temporo-parietal electrodes where the cooperative situation elicited more positive amplitudes. Further, the differences induced by the social situations were stronger in participants with higher scores on a perspective taking test. In sum, our results replicate previous studies about the FRN and extend them by comparing neurophysiological responses to positive and negative outcomes in a task that simultaneously engages two participants in competitive and cooperative situations.

## Introduction

In every day life, humans frequently commit errors. For example, they are prone to press incorrect buttons, trip over household objects or make typing mistakes. These errors often influence not only the person committing the mistake but also other people. Such erroneous actions may have a negative impact on others if people are cooperating in a task (e.g., moving furniture together). Conversely, they may have a positive impact on others if people are competing in a task (e.g., in a game of table tennis). These mistakes that involve others frequently require external feedback to find out about the impact of one’s own and others’ performed actions. Thus, it is likely that the human brain has mechanisms that distinguish between positive and negative outcomes of one’s own and others’ actions.

Earlier research on error processing in tasks performed individually shows that humans have a fast and efficient error detection mechanism (1, 2). In particular, studies using electroencephalography (EEG) identified event-related-potential (ERP) components instantly following one’s own errors awareness, or feedback regarding the outcome of one’s own actions (3). These components are known as error-related-negativity (ERN) and feedback-related-negativity (FRN). The ERN is evoked 50-70 milliseconds after an erroneous action is carried out (e.g., an incorrect button press) and it originates from the anterior cingulate cortex and the presupplementary motor area in the posterior medial frontal cortex (4–6). The FRN is elicited approximately 200-350 milliseconds after performance feedback is received and is considered to have a similar origin as the ERN (7). Holroyd and Coles (2002) proposed that the ERN/FRN component is elicited as soon as the outcome of an action can be detected by proprioceptive, motor or external feedback. They also proposed a direct relationship between a negative outcome detection and reward processing. In essence, whenever the result of an action is worse than expected, which results in a loss of reward, the ERN/FRN is elicited.

While these components have been widely studied in individuals, little research has investigated how humans process feedback about actions that involve others. A first step in this direction was made by van Schie, Mars, Coles & Bekkering (2004). They found that the FRN component occurs after observing an error committed by others. Given the sensitivity of the FRN to mistakes of others, researchers suggest that it might reflect the processing of socially relevant stimuli. Further studies explored this idea by manipulating the social situation (i.e., either cooperative or competitive) while participants performed or observed actions and received feedback about monetary rewards (8, 9). Results showed that the FRN was elicited by losses of others in a cooperative situation. In a competitive situation, conversely, others’ gains elicited the FRN. These results indicate that the FRN reflects the valence of an outcome, which in turn depends on the current social situation.

In contrast to studies of the FRN discussed above, the ERN, which is elicited for self-generated errors, appears to be not influenced by the social situation (10). In another study self-generated errors elicited ERN in both cooperative and com-petitive situations, however, observed errors of others elicited the FRN only in a cooperative situation (11). These studies focused on outcome processing in cooperative and competitive situations. However, the tasks used in these studies involved actions that are performed in turns and there was always either a division between a performer and observer participant (9–11) or the partner was virtual (8). Hence, it is not clear whether these findings would also generalize to designs in which co-actors perform a task jointly.

To close this gap in the literature, set of recent studies also investigated the FRN in situations in which humans perform tasks together. Humans in real life often perform actions together with others, instead of observing another human performing an action alone. Thus, studying the social aspect of outcome processing requires paradigms, in which co-actor performs tasks jointly (12, 13). In line with this idea Picton, Saunders and Jentzsch (2012) tested dyads of participants in a cooperative joint choice reaction time task. In their study, participants were able to realize their own mistakes without feedback, which elicited the ERN, while mistakes of a partner had to be inferred from visual feedback, which elicited the FRN (14). In an even more naturalistic set-up, Loehr and colleagues tested piano duets (15, 16). Such a music paradigm allowed for a clear division between one’s own, other’s and joint errors. Results of both Picton et al’s and Loehr et al’s experiments confirmed that the FRN monitors both one’s own and other’s errors in joint situations. Interestingly, the FRN is stronger for one’s own than joint mistakes, and stronger for joint mistakes than others’ mistakes (16). These studies focused on the monitoring of actions in cooperative joint set-ups. However, according to our knowledge there are no studies that involve two participants performing actions and receiving feedback about their individual and joint actions in both cooperative and competitive situations.

To fill this gap in the literature, in the present study we focused on two aspects: First, in our experiment both participants were actively performing a task. That is, in contrast to previous research there was no distinction between an active co-actor and a passively observing co-actor (8, 9). Instead, each of the participants performed their individually assigned task in parallel and observed their own and the co-actor’s errors. Second, rewards (positive, negative and neutral) associated with errors depended on whether the assigned task was performed in a cooperative or competitive situation. With this design, the main question we addressed was whether the FRN is influenced b y d ifferent s ocial s ituations w hen both co-actors actively perform a task. Additionally, by including neutral conditions (i.e., condition without any monetary rewards) in the design, we were able to investigate whether FRN amplitudes differed between errors that are associated with monetary outcomes (positive and negative) and errors that are not associated with any monetary rewards (neutral). Such comparisons were only rarely addressed in previous research (17). We also aimed to relate FRN amplitudes to personality traits measured with a questionnaire. Namely, we focused on the Perspective taking subscale of the Interpersonal reactivity index (IRI, (18)) that measures the tendency to spontaneously adopt the psychological point of view of others. We chose this subscale because it was already shown that FRN amplitudes correlate with the Perspective taking scores (19). Finally, we performed exploratory analysis to explore the time course of processing feedback about self-produced actions and co-actors’ actions depending on the social situation.

## Methods

### Participants

52 students (35 females, mean age = 24.1, standard deviation = 4 years) randomly grouped into in 26 dyads participated in the experiment. Prior to the experiment we asked all participants whether they knew each other and paired only strangers. The ethics committee of the University of Osnabrück approved the experiment. We informed participants about their rights and all participants signed a written consent form. Participants could chose either a monetary reward or course credits in exchange for their participation. All participants where we measured EEG opted for the monetary reward.

### General Apparatus

We tested participants in dyads. They sat next to each other on the same side of a table in the same room. To avoid interference and communication during the experiment, we separated them with a cardboard screen (Figure 1, B). We presented stimuli on two identical computer monitors (BenQ 24 inches, 1920×1080 pixels, refresh rate 120 Hz). We used two separate keyboards (Cherry RS 6000) to collect behavioral responses, one for each participant. The experiment was programmed using the Python library PsychoPy (20) and the experimental procedure and data collection were implemented in Python 2.7.3. The experiment was run on an Intel Xeon computer.

**Fig. 1.**
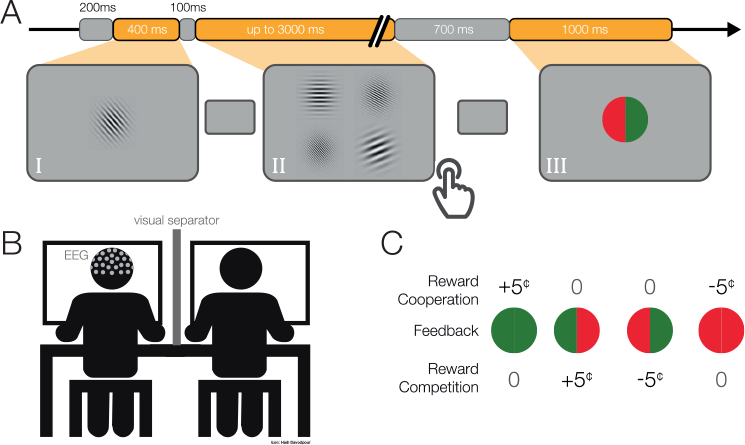
A) Single-trial. We presented a gray mask for 200 ms, followed by (I) a target Gabor patch for 400 ms. Then again, a gray mask was displayed for 100 ms, followed by (II) four Gabor patches until both participants responded (maximum 3000 ms). Subsequently, we displayed a gray mask for 700-800 ms, followed by (III) the feedback for 1000 ms. B) Schematic depiction of the experimental setup. C) Pay-off matrix. Participants received rewards differently in cooperative and competitive situations.

### Experimental design

Each member of a dyad performed a four-alternative forced-choice (4-AFC) visual task (Figure 1, A) and later received feedback about their performance and associated monetary rewards (Figure 1, C). In each dyad, one participant performed an orientation discrimination task and the other participant a spatial frequency discrimination task. First, we presented a target object in the middle of the screen for 400 milliseconds. The target object was a single Gabor Patch of size 9.95°×9.95° visual angle, oriented at a randomly chosen angle (between 20°−80° and 100°-160°) and with a randomly chosen spatial frequency (between 10 and 20 cycles/stimulus size). Subsequently, we displayed a gray mask with a fixation cross in the middle (linewidth of 0.13° visual angle) for 100 milliseconds followed by four Gabor patches arranged in a 2×2 grid, each patch separated from neighboring patches by 0.41° visual angle on each side. Each of the four Gabor patches was of the same size as the target object. One Gabor Patch always had the same orientation as the target object while the other three patches were manipulated according to a QUEST staircase procedure (21). A different Gabor patch had the same spatial frequency as the target object and the other three patches again had different spatial frequencies according to a second QUEST staircase procedure (for more details about the QUEST procedure, see Experimental procedure). The location of the correct answer for each of the participants was randomized between four possible locations. Participants responded with key presses (‘Q’,‘W’,‘A’,‘S’or ‘7’,‘8’,‘4’,5’ on the num-pad, for the participants seated on the left or right respectively). The key corresponded spatially with the displayed Gabor patches. We displayed Gabor patches until both of the participants gave their responses or 3000 milliseconds passed. In the case of no response, the answer was considered as incorrect. We instructed participants to give their answers as accurately and as quickly as possible. Subsequently, a gray mask with a fixation c ross was d isplayed f or 700-800 milliseconds and then feedback appeared on the screen. We used a colored circle (radius: 3.94° visual angle) vertically divided in halves to inform participants about the performance of both participants. The color of the feedback was dependent on the participants’ answers. The green color indicated correct answers and red incorrect answers. The left semicircle and right semicircle gave feedback to the left and right participants, respectively. Additionally, we presented individually a letter (0.8° visual angle, ‘W’ for wins, ‘L’ for losses and ‘T’ for ties; for more details, see subsection “Social manipulation and monetary rewards” below) in the middle of a circle. Feedback was displayed for 1000 milliseconds and was followed by a gray mask for 200 milliseconds before moving on to the next trial (Figure 1, A).

### Social manipulation and monetary rewards

The feedback included information about individual and joint errors as well as the resulting positive, negative or neutral monetary rewards. Note, the schema of monetary rewards, as given in the pay-off matrix, defined the social situation as cooperative or competitive. The gain or loss of 5 cents was dependent on the particular social situation as follows (Figure 1, C):

In the cooperative situation the trial was considered as a win, and consequently positively rewarded, only in the case in which both of the participants responded correctly (one green semi-circle for each of the two participants). In the case that both participants were wrong, it was considered a loss and as a negative reward five cents were subtracted from their budgets (one red semi-circle for each of the two participants). In the case that one participant was correct and the other was incorrect, no money was added to or subtracted from either budget (half green and half red circle).

In the competitive situation both participants answering correctly or incorrectly resulted in a tie (full green or red circle). Thus, no money was added to or subtracted from either budget. A reward was achieved when one participant was correct and the other was incorrect (half green and half red circle). In this case the reward was added to the correct participant’s budget and subtracted from the incorrect participant’s budget. At the end of each block the participants’ respective budgets were calculated and displayed on the screen.

Social situations alternated between blocks. At the beginning of each block, we provided information regarding the block number, the social situation and rewards associated with each feedback. In addition, “win” or “lose” was shown as text inside the feedback stimulus (Figure 1, C). Each participant had an initial budget of 10 Euro that could increase or decrease by 5 cents based on their performances in each trial.

### Experimental procedure

One participant of each dyad was prepared for the EEG recordings outside of the recording chamber. After around 45 minutes, when preparation was finished, both participants were seated side-by-side in a room at a 60 cm distance to their screen. For technical reasons, the participant measured with the EEG sat on the left side. The experimental session lasted approximately 90 minutes and wdure (21) was performed for each participant foras structured as follows: After detailed written and oral instructions, a QUEST staircase proce the assigned task with the goal to home in on 50% performance, i.e. well above the chance level of 25%. To achieve this, we used the PsychoPy QuestHandler function with the threshold set to .63 and a gamma .01. Both participants performed 100 training trials. For the participant performing the orientation discrimination task we varied the degree of orientation between 1° and 45° with a starting value of 15° and a standard deviation 10°. For the other participant, who performed the spatial frequency discrimination task, we varied the spatial frequency between 1 and 25 cycles/stimulus size with starting value of 3 cycles/stimulus with a standard deviation of 3 cycles/stimulus. Subsequently, participants proceeded to the actual experiment, which consisted of a total of 640 trials grouped in 16 blocks of 40 trials each. After 20 trials in each block, participants were asked to answer in which social situation they were currently in. Namely, they were asked to indicate whether the current block was a cooperative or competitive situation, in order to check whether the participants remembered the social situation manipulation correctly. Blocks were separated by short rests and the overall experiment was divided into three parts with short breaks. In these breaks experimenters made sure that participants were not exchanging any information about the experiment. When the tasks were completed, participants filled out the Interpersonal reactivity index (IRI, 28 questions) questionnaire (18).

### Methods of EEG data acquisition and preprocessing

Electrophysiological data were recorded using a 64-Ag/ AgCl electrode system (ANT Neuro, Enschede, Netherlands), using a REFA-2 amplifier (TMSi, Enschede, Netherlands) with electrodes placed on a Waveguard cap according to the 5% electrode system ((22)). The data was recorded using average reference electrode at a sampling rate of 1024 Hz. Impedances of all electrodes were manually checked to be below 10 kΩ before each experiment. We used R and MATLAB to preprocess and analyze the data. All analysis scripts and data are available online (https://osf.io/c4wkx/). We used the eegvis toolbox (23) to visualize the exploratory analyses. Data were preprocessed using the EEGLAB toolbox (24) in the following order: First, the data were downsampled to 512 Hz and subsequently filtered using a 0.1 Hz high-pass filter and a 120 Hz low pass filter (6 dB cutoff at 0.5Hz, 1 Hz transition bandwidth, FIRFILT, EEGLAB plugin). Channels exhibiting either excessive noise or strong drifts were manually detected and removed (2.1 +/− 2.5, mean and standard deviation respectively). After this, the continuous data were manually cleaned, rejecting data sequences including jumps, muscle artifacts, and other sources of noise. To remove eye and muscle movement-related artifacts, an independent component analysis based on the AMICA algorithm (25) was computed on the cleaned data. The independent components (ICs) corresponding to eye, heart, or muscle activity were manually selected based on their timecourse, spectra and topography, and removed before transforming the data back into the original sensor space (number of removed ICs 8.3 +/− 5.2, mean and standard deviation, respectively). The initially removed channels were interpolated based on the activity of their neighboring channels (spherical interpolation). Subsequently, the continuous data were divided into epochs for each trial by including data from 200 millisecond pre-stimulus to 1000 millisecond post stimulus, using the time window between −200 millisecond and stimulus onset for baseline correction. For the exploratory analysis we used 62 electrodes (Fp1, FPz, Fp2, F7, F3, Fz, F4, F8, FC5, FC1, FC2, FC6, T7, C3, Cz, C4, T8, CP5, CP1, CP2, CP6, P7, P3, Pz, P4, P8, POz, O1, Oz, O2, AF7, AF3, AF4, AF8, F5, F1, F2, F6, FC3, FCz, FC4, C5, C1, C2, C6, CP3, CPz, CP4, P5, P1, P2, P6, PO5, PO3, PO4, PO6, FT7, FT8, TP7, TP8, O7, PO8).

## Results

### Behavioral analysis

#### Social situation awareness

To assure that participants payed attention to the different social situations in the experiment we asked them in the middle of each block whether the current block was a cooperative or competitive situation. Answering this question participants achieved a high accuracy (mean correct answers = 97%, standard deviation = 7%), suggesting that participants consistently understood and memorized the instructions about differences between social situations.

#### Accuracy

Prior to running the actual experiment, we used a QUEST staircase procedure to adjust a difficulty in each task for participants such that participants were expected to attain a 50% accuracy. Confirming this expectation, the mean accuracy in the task was 53% (standard deviation = 9%) and the mean difference between paired participants was 8% (standard deviation = 6%). It was important that both paired participants performed with comparable accuracy to avoid that the analyzed ERPs are influenced by differences at the behavioral level. Further, it results in an even distribution of performance data in correct-correct, correct-false, false-correct and false-false.

#### Response time

We analyzed response times to test whether our experimental manipulations influenced behavioral responses. Prior to analysis, we excluded all trials with response times faster than 50 milliseconds (2 trials) because such fast responses are likely due to premature responses. Then, we used a linear mixed model (LMM) to analyze response times. The LMM was calculated with the lme4 package (26) and *p*-values were based on Walds-T test using the lmerTest package. Degrees of freedoms were calculated using the Satterthwaite approximation. We modeled responses times by task, social situation and correctness as fixed effects and interactions between them. As random effects, we used random intercepts for grouping variables participants and dyads. In addition, we used random slopes for all fixed effects, including interactions, in the participant grouping variable. For all predictors, we used an effect coding scheme with binary factors coded as −0.5 and 0.5. Thus the resulting estimates can be directly interpreted as the main effects. The advantage of this coding scheme is that the fixed effect intercept is estimated as the grand average across all conditions and not the average of the baseline condition. We found a main effect of correctness (*t*(50.18)= −8.1, *p* <.0001). Correct answers were on average 80 milliseconds faster than incorrect answers. The main effects for the two other predictors (tasks and social situations) and all possible interactions were not significant (*p* >.17). These results suggest that different tasks (orientation and spatial frequency) and social situations (cooperative and competitive) are of comparable level of difficulty and engage two participants to similar degrees.

### Electrophysiological data

To analyze EEG data in form of ERPs, we applied a prese-lected single-trial based LMM analysis (27). We defined the FRN as the mean amplitude over six electrodes (Fz, F1, F2, FCz, FC1, FC2) between 200 and 300 milliseconds after the feedback of each trial. Our choice of electrodes and time window was based on previous research and were pre-specified before any analysis (28). We modeled the FRN using outcomes (win and lose) and social situations (cooperative and competitive) as fixed effects and an interaction between them. As random effects, we modeled random intercepts for participants and random slopes for both predictors (outcomes and social situations) and interaction between them. For the same reason as above, predictors were effect coded, i.e., binary factors are coded as −0.5 and 0.5. The result of this analysis are presented in Table 1 and ERPs in Figure 2. We found main effects for the outcome (*t*(26.02) = −5.85, *p* <.001) and the social situation (*t*(26.01) = 4.4, *p* <.001). The interaction between these factors was not significant (*t*(27.15) = −.93, *p* = .36). These results suggest that the FRN differs between positive and negative outcomes and between cooperative and competitive social situations and that these two effects are independent of each other.

**Table 1.**
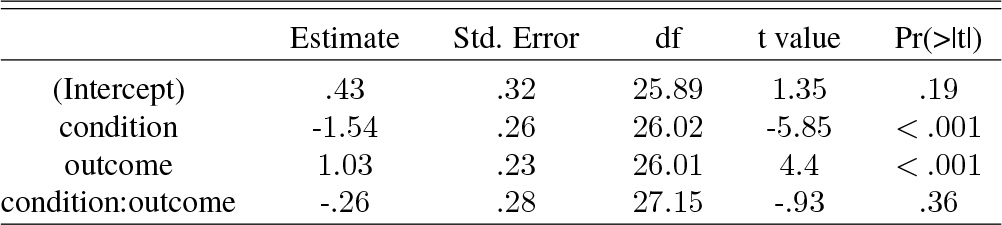
LMM Effects of outcome and social situation on the FRN (mean amplitude (200-300 milliseconds)) (effect coding: −0.5,0.5, maximal LMM).

**Fig. 2.**
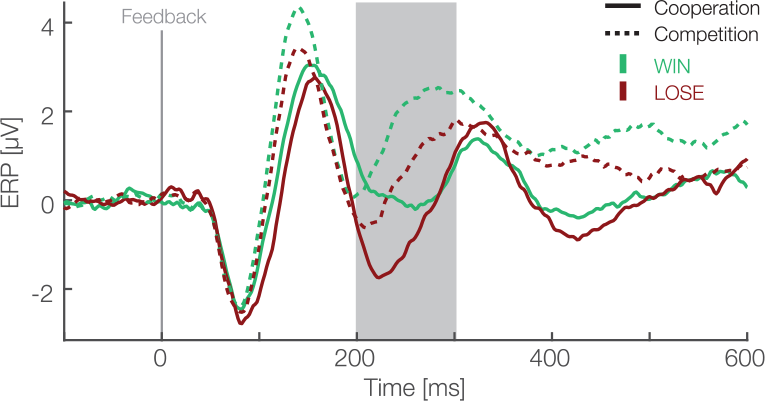
Feedback locked ERP waveforms at pooled electrode sites (F1, Fz, F2, FC1, FCz, FC2). Data are averaged referenced. Green and red colors represent the outcome, i.e., win and lose trials respectively. Solid and dashed lines represent cooperative and competitive situations. The gray box shows the preselected time window used for the confirmatory statistical analysis (200-300 ms).

Additionally, we used individual estimates of the difference between the FRN in the two social situations to correlate them with the Perspective Taking Score. We calculated the Spearman’s Rho to quantify the association of the Perspective taking score and individual participant’s mixed model best linear unbiased prediction of the factor social situation from the mean amplitude analysis. We chose Spearman’s correlation because our questionnaire data was rank data. We found a significant negative correlation (*r* = −.54, *p* = .005, Figure 3). This result suggests that on average the effect of the social situation is stronger on the characteristic ERPs in participants with personality traits related to high perspective taking abilities.

**Fig. 3.**
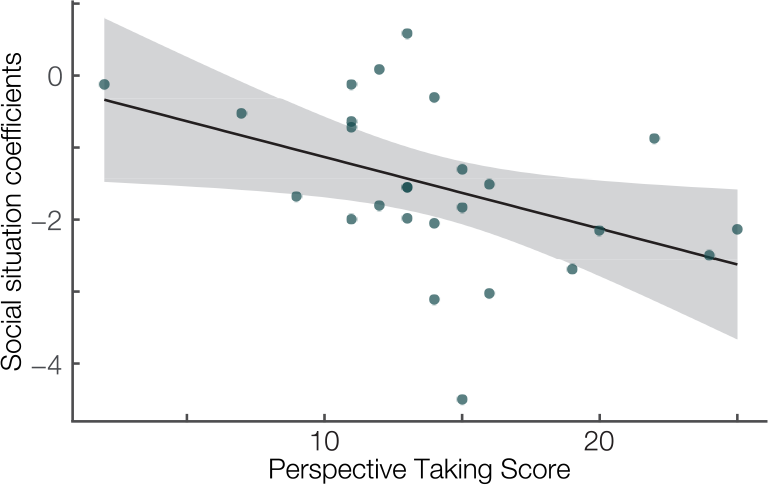
Correlation between the Perspective taking subscale of the Interpersonal Reactivity Index (Davis, 1983) (x-axis) and the single-subject linear mix model estimates of the social situation effect (y-axis). Spearman’s *r* = −.54, *p* = .005. Linear fit with 95% confidence interval.

Furthermore, after visual inspection of the grand average ERPs (Figure 2), we decided to also apply a peak to peak amplitude analysis because the FRN peaked earlier than expected (29). For the peak to peak analysis we used the same electrodes as for the mean amplitude analysis (Fz, F1, F2, FCz, FC1, FC2). We used the grand average to identify the maximum positive peak between 140 and 200 milliseconds and the maximum negative peak between 200 and 270 milliseconds after feedback presentation. We subtracted the average maximum negative peak amplitude from the average maximum positive peak over these time windows. Then, we applied exactly the same LMM analysis as with the mean amplitude (details above). The result of this analysis are presented in the Table 2. We found main effects for the outcome (*t*(26) = 3.55, *p* = .001) and the social situation (*t*(26.09) = −3.04, *p* = .005). The interaction between these factors was not significant (*t*(25.98) = −.98, *p* = .34). These results are in line with results of mean amplitude analysis, further corroborating that FRN amplitudes differ between positive and negative outcomes and between cooperative and competitive situations.

**Table 2.** Effects of outcome and social situation on the FRN (peak to peak amplitude (140-270 milliseconds)) (effect coding: −0.5,0.5, maximal LMM).

|  | Estimate | Std. Error | df | t value | Pr(>\|t\|) |
| --- | --- | --- | --- | --- | --- |
| (Intercept) | 12.86 | .6 | 25.99 | 21.55 | 0 |
| condition | 1.3 | .36 | 26 | 3.55 | .001 |
| outcome | -.71 | .23 | 26.09 | -3.04 | .005 |
| condition:outcome | -.41 | .42 | 25.98 | -.98 | .34 |

Next, we analyzed differences in the FRN amplitudes between monetary versus neutral outcomes crossed with social situations. For this, we utilized a difference wave approach (30). In each of the social situations, we subtracted ERPs of negative from positive monetary outcomes. In addition we subtracted incorrect responses from correct ones in the neutral monetary outcomes. Then, we quantified the FRN as the mean amplitude between 200 and 300 milliseconds after the feedback presentation for each condition. We used a two-way repeated measures ANOVA with social situation (cooperative vs. competitive) and type of outcome (monetary vs. neutral) as within participant factors. We found a main effect of social situation (*F*(1,25) = 6.17, *p* = .02, *η*^2^ = .022, Figure 4), a main effect of type of outcome (*F*(1,25) = 4.55, *p* = .04, *η*^2^ = .026) and no interaction between these factors (*F*(1,25) = .34, *p* = .56, *η*^2^ = .003). These results suggest that the effect of the social situation on the FRN reported above extends to neutral outcomes. Furthermore, the significant difference between monetary and neutral outcomes suggest that is sensitive to both monetary rewards and task performance.

**Fig. 4.**
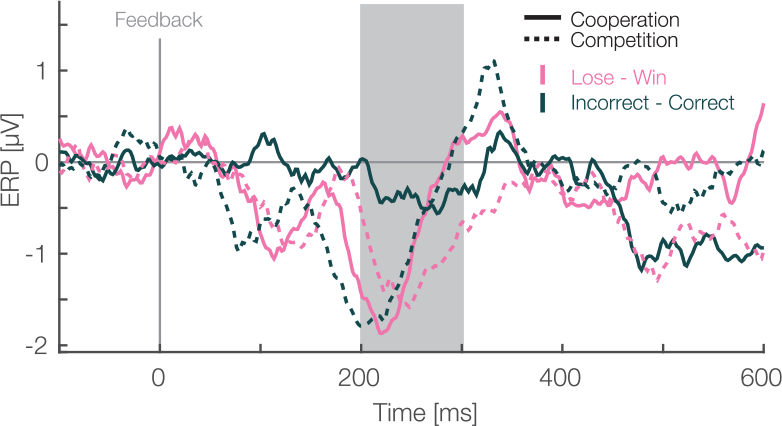
Feedback locked difference waveforms at pooled electrode sites (F1, Fz, F2, FC1, FCz, FC2). Data are average referenced. Pink and green colors represent the monetary outcome, i.e., lose-win and incorrect-correct trials respectively. Solid and dashed lines represent cooperative and competitive situations. The gray box shows the preselected time window used for the confirmatory statistical analysis (200-300 ms).

For the exploratory analysis, we used the Threshold-FreeCluster-Enhancement method (TFCE) and permutation analysis (31–33). This method allows for comparisons between experimental conditions over all electrodes and time points of ERPs while at the same time controlling for the multiple comparison. We analyzed the EEG data with a two-way repeated measures ANOVA with outcome (win vs. lose) and the social situation (cooperative vs. competitive) as within participants factors and taking into account 62 electrodes, and all time points between 0 and 600 milliseconds. We enhanced the signal with the TFCE method and used permutation tests to account for multiple comparisons. We used 5000 permutations and for each permutation we randomized the assignment to different experimental conditions of each data point within each participant. For each of these TFCE permutations, a repeated measures ANOVA was calculated. The maximum F-value across chosen samples in time and space were used to construct a max F-value distribution, against which the actual F-values were compared. We considered F-values above the 95th percentile to be significant. The results of this analysis are presented in Figure 5. We found two separate clusters of significant a ctivity f or t he m ain e ffect o f o utcome. One cluster spans from 88 to 152 milliseconds (median *p* value: *p* = .01, min *p* value: *p* = .001) with a peak at C1 electrode 121 milliseconds after the feedback. The other cluster ranges between 172 and 340 milliseconds with a peak at Fz electrode 240 milliseconds following the feedback (median *p* value: *p* = .01, min *p* value = .0006). This cluster resembles spatially and temporally the FRN and it was more negative for lose than win outcomes. Moreover, we found that there is a main effect of the social situation. This cluster stretched from 68 till 600 milliseconds (median *p* value: *p* = .0004, min *p* value: *p* = .0002) and encompassed all electrodes at different time points, suggesting a robust difference in processing of feedback between cooperative and competitive situations. The peak significant value was at FC5 electrode 143 milliseconds after the feedback. Overall, these results support the observations above of large differences between processing of feedback between cooperative and competitive situations and suggest that the difference in processing positive and negative feedback starts earlier than classically considered time window for the FRN.

**Fig. 5.**
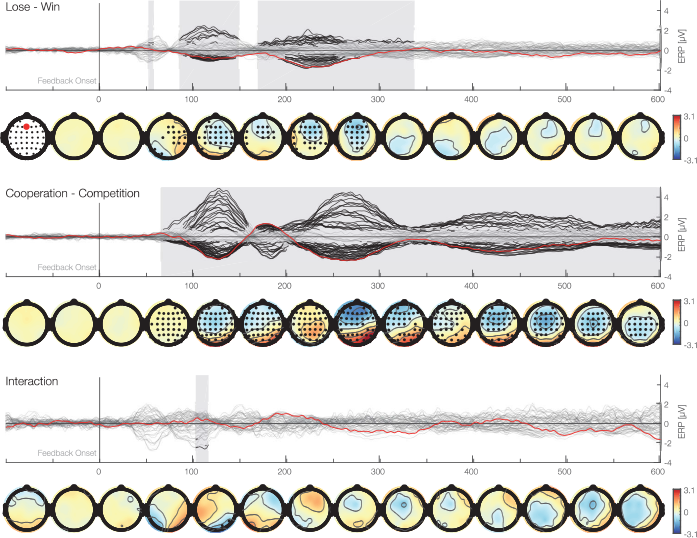
Time-series plots of the EEG amplitudes of the main factors and interaction for each electrode aligned to the feedback stimulus. First row (butterfly plot) shows time against activity of all electrodes. In addition, the Fz electrode is marked in red. Black marked clusters are significant under a TFCE permutation p-value of TFCE corrects for multiple comparisons over time and electrodes. Second row shows topographical plots representing the mean amplitudes averaged over 50 ms bins. Black marked electrodes represent significant channels.

Lastly, we address a potential visual confound in our design. As we used four different visual stimuli to inform our participants about their performance and associated rewards, results potentially reflect differences of the visual feedback. To address this potential perceptual confound, we invited five participants again, who previously completed the experiment, for a control experiment. In this version of the experiment, the Gabor patches were not displayed and random feedback was provided. Thus, this experiment controls for the pure visual effect of the feedback. To assess this potential confound, we calculated grand average ERPs for experimental and control data. We visually inspected the ERPs and found no difference between the different visual feedback displays, including the early visual components. Then, we subtracted these control data from the experimental data. Again, visual inspection suggests that differences in the visual appearance of the feedback information did not influence the FRN (Figure 6). This is in line with previous research that shows only early components e.g. C1, P1, N1, in the first 150 milliseconds are modulated by such low-level visual stimuli properties (34, 35). Hence,we are reassured that our results represent differences between outcomes and social situations and not due to differences in the visual stimuli.

**Fig. 6.**
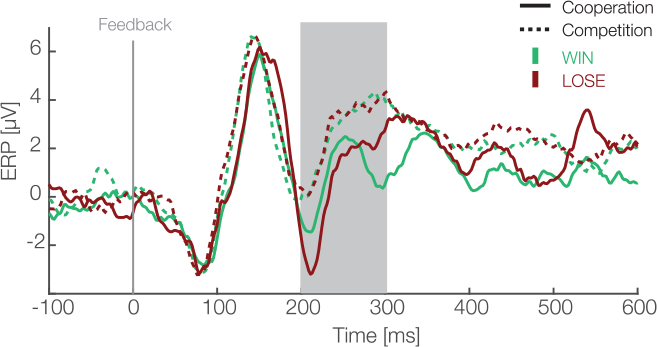
Feedback locked averaged difference waves between experimental and control data for 5 participants pooled at electrode sites (F1, Fz, F2, FC1, FCz, FC2) are shown. In experimental data green and red colors represent the outcome, i.e., win and lose trials respectively, while solid and dashed lines represent cooperative and competitive situations. Difference between social situations and outcomes is visible after subtraction of electrophysiological response to identical visual stimuli without any content. This suggests that our results represent differences in experimental manipulations but not visual properties of stimuli.

## Discussion

The goal of the present study was to compare reward processing between different social situations as well as to test whether earlier results (14) generalize to a setting which actively involves two participants. For this purpose, we designed a joint 4-AFC visual task, in which two co-actors both concurrently perform a task and receive rewards depending on the social situation. We were able to replicate the difference in FRN amplitudes between positive and negative outcomes in the cooperative situation (14). Moreover, we extended these earlier results by observing a significant difference between win and lose outcomes in the competitive situation. We also found that the FRN significantly differs between social situations, suggesting that reward processing is modulated by the social situation. Further, the difference induced by the social situations were stronger in participants with higher perspective taking scores, which were obtained using a perspective taking questionnaire. Finally, we compared feedbacks with and without monetary outcomes (win/lose vs. neutral) in both social situations. We found that our reported effect, that the social situation affects the FRN, also extends to the processing of neutral outcomes. Moreover, we found a significant d ifference b etween feedbacks with and without monetary outcomes, suggesting that the FRN is sensitive to both monetary rewards and task performance.

Earlier behavioral findings support the idea that humans co-represent co-actors actions even if they are irrelevant to one’s own goals (36), for a recent general review, see: (37). Such representations may also influence how humans process feedback about actions and associated monetary rewards while performing joint actions with another person. Therefore, our experiment involved two participants performing their tasks simultaneously and hence differs from previous studies that utilized a virtual partner to investigate differences between social situations (8). Moreover, the design allows for concurrent actions from both participants – an aspect that it is not present in designs that employ turn-taking tasks which create a division between a performer and observer (9–11). Thus, with the results of the present study, we extended earlier findings by demonstrating that they also generalize to a setting involving co-actors that both actively and simultaneously perform a task.

Our result that the outcome (positive vs. negative reward) affects the FRN in both social situations is in line with a great body of earlier research(28). We quantified the FRN in two different ways (mean and peak to peak amplitude) and applied additional exploratory analyses. Results of all three analyses provide strong evidence that negative outcomes elicit more negative amplitudes at mid-line electrodes around 200 to 300 milliseconds after the feedback presentation. Such an outcome of our study suggests that the FRN component is robust and it generalizes from individual to joint set-ups and different social situations. In contrast, our results are not compatible with the theory that the FRN represents differences in expectancies and probabilities (38, 39). In our task the probabilities for each outcome were nearly equal, therefore, there are no differences in probabilities or expectancies. Future studies could investigate whether reward processing is also affected by the outcome in tasks, in which both co-actors actively perform a task collaboratively as, for instance, in joint perceptual tasks (e.g., (40); (41); (42); (43); (44); for a recent review, see (45)) or in joint motor tasks (e.g., (46), for a recent review, see (47)).

The main question, namely, whether reward processing differs between social situations was addressed in three ways. First, we analyzed the FRN as mean as well as peak to peak amplitude and found a main effect of social situation. Second, we also found a main effect of social situation when analyzing the difference waves. Third, using an exploratory analysis, we again found a main effect of social situation. Taken together, these results suggest that the FRN amplitudes are affected by the social situation. This raises the question which aspect of the change in social situation affects the FRN. Potentially, the social situations might differ with respect to arousal state and the amount of attentional resources utilized. However, we did not observe differences in the level of performance as a function of the social situation. This makes an influence on the FRN by variations of arousal or attentional resources unlikely. Therefore, our study provides evidence that reward processing is affected by social situations, however, further research is needed to unravel details of involved processes.

A previous study suggested that that the FRN is only sensitive to the outcome, but not task performance as such (8). As studying the FRN in response to neutral outcomes is mostly neglected in literature (but see (17)), this is difficult to disentangle. Due to our design that included neutral outcomes, we were in a better position. Specifically, the comparison of FRN amplitudes between feedbacks with and without monetary outcomes in combination with correct or incorrect individual performance, results in a significant difference between feedbacks with and without monetary outcomes. This result suggests that different neural processes are involved in processing outcomes and task performance. Given that we find that the FRN is present for neutral outcomes, this result suggests that the FRN is sensitive to outcome as well as task performance. They suggested that the FRN is only sensitive to the outcome.

In this study, we used state of the art EEG analysis methods, namely Linear Mixed Models for hierarchical analysis of single trial activity (27) and TFCE to control for multiple comparisons (31, 32). In the following, we first provide a discussion of the benefits using these analysis techniques and then further discuss the obtained results of our exploratory analysis. We quantified the FRN on a single trial basis and used the LMM to model the FRN. This approach helps to account for a multitude of problems. For instance it handles unequal number of observations per cell, allows for between participant variability in effect sizes and combines single participant variability and group level variability (48–51). In our experiment, we tried to reduce the first p roblem o f u nequal cell size by using the QUEST procedure to obtain almost equal number of trials. Nevertheless, EEG data has to be cleaned and depending on the noise level the number of rejected trials varies between participants. However, the issue of high variability between participants in cognitive neuroscience field is prevalent and has to be accounted for (52). The LMM approach is suitable to address this problem. Our additional motivation to use this method was related to its capability of estimating effect sizes for individual participants. We used those to correlate them with information about personality traits of participants to test a possible association between neurophysiological and questionnaire data. We also made use of the TFCE permutation analysis to perform the exploratory analysis (32) without specifying electrode sites or time window. This approach circumvents the need to preselect time points and electrodes (53), which is an additional benefit as making these decisions may not always be straightforward, especially in the absence of clear guidelines.

Using this exploratory analysis, we found the same pattern of results as above in our confirmatory analysis. Namely, we found a main effect of the outcome and social situation in both the LMM and the permutation analysis, further corroborating earlier results that the FRN is sensitive to positive and negative outcomes and the social situation (28). In addition, our exploratory analysis showed that these differences for the FRN preceded the time window typically defined for the FRN, suggesting that the human brain differentiates the valence of the outcome and the social situation earlier than previously suggested (9–11, 14–16, 54). Our results (Figure 5, second row), suggest that there are stronger positive activation in cooperative than competitive situation in two stages of processing of the feedback. Namely, around 160 and 280 milliseconds after the feedback presentation. The social situation main effect might arise from a source close to CP6 and P6 electrode. Because Superior temporal sulcus (STS) and Temporoparietal junction (TPJ) are close to these electrodes and earlier fMRI research suggests these areas are involved in differentiating the self from others, this might be the origin (55)Ṫ hus, it might be interpreted that while people receive feedback and process them simultaneously in a coop-erative situation they merge their own and their co-actor positive outcomes and process them as simultaneously while the competitive situation requires distinct processing of rewards. However, this interpretation have to be taken cautiously due to the inverse problem.

Moreover, we investigated the relation between the Perspective taking score and mixed model best linear unbiased prediction of the factor social situation. We found that the higher the Perspective taking score, the stronger is the difference in FRN amplitudes between social situations. This result suggests that personality traits related to perceiving and understanding others might be related to the strength of the neurophysiological response to rewards. Thus, brain mechanisms involved in reward processing in people showing more consideration for others, might be more sensitive for different social situations. However, this result and interpretation should be treated with caution, as using mixed model best linear unbiased prediction in combination with a correlation analysis is a new approach and still has to be fully validated (56).

Taken together, we investigated neural underpinnings of feedback processing in cooperative and competitive situations. We find that the FRN component is sensitive not only to positive and negative outcomes but also to the social situation in a design, in which both co-actors in dyad actively perform a task.

**Fig. 7.**
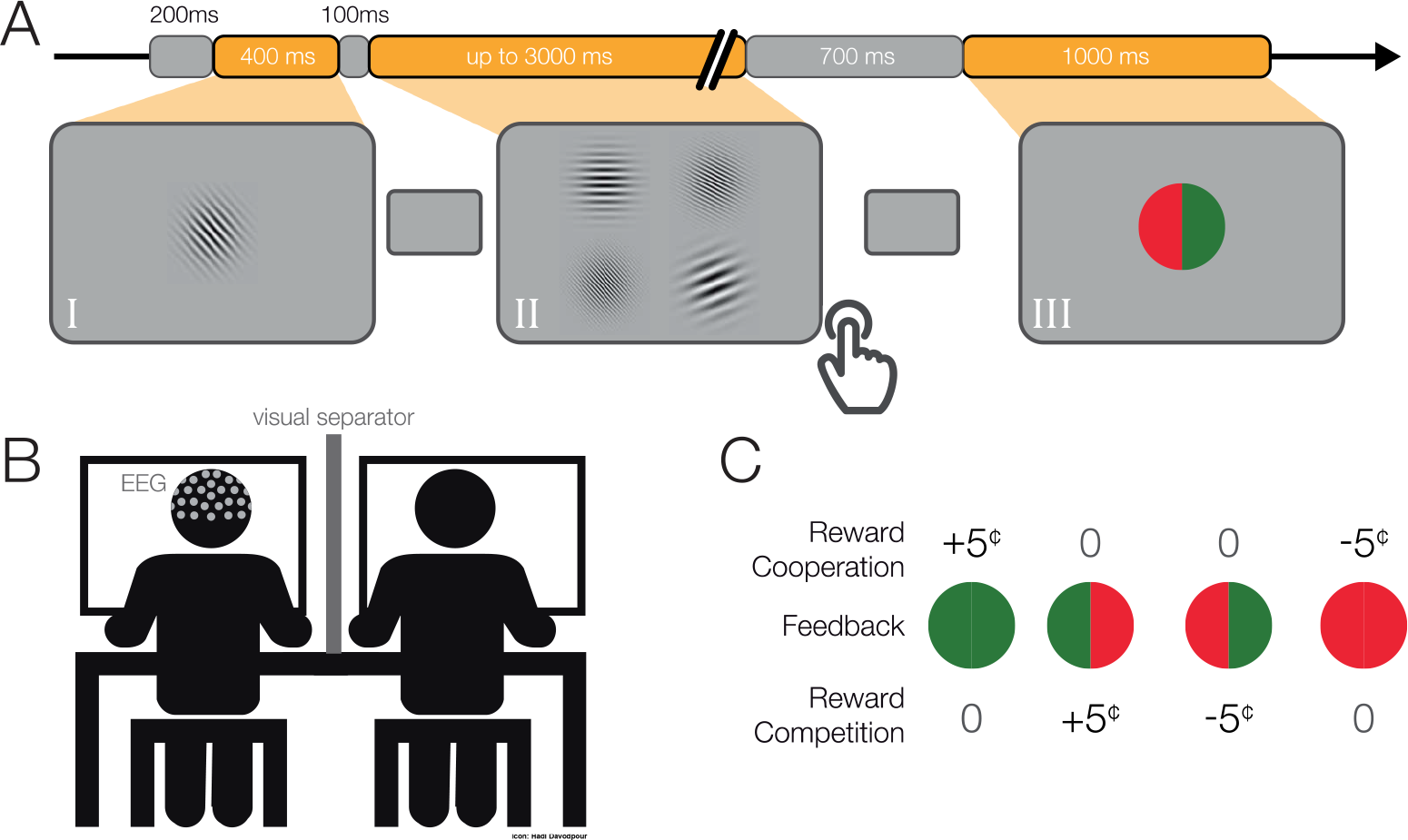
A) Single-trial. We presented a gray mask for 200 ms, followed by (I) a target Gabor patch for 400 ms. Then again, a gray mask was displayed for 100 ms, followed by (II) four Gabor patches until both participants responded (maximum 3000 ms). Subsequently, we displayed a gray mask for 700-800 ms, followed by (III) the feedback for 1000 ms. B) Schematic depiction of the experimental set-up. C) Pay-off matrix. Participants received rewards differently in cooperative and competitive situations.

**Fig. 8.**
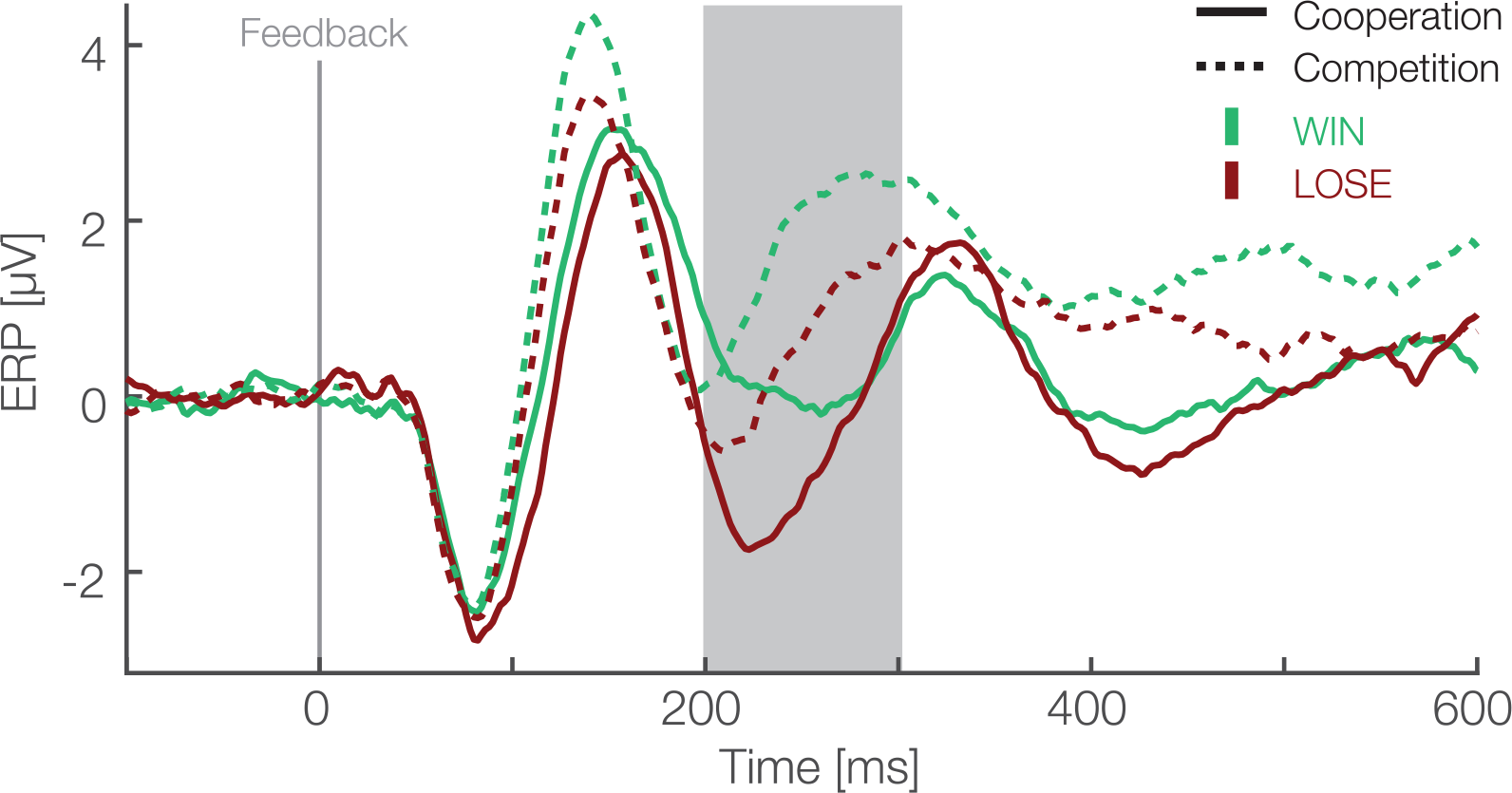
Feedback locked ERP waveforms at pooled electrode sites (F1, Fz, F2, FC1, FCz, FC2). Data are averaged referenced. Green and red colors represent the outcome, i.e., win and lose trials respectively. Solid and dashed lines represent cooperative and competitive situations. The gray box shows the preselected time window used for the confirmatory statistical analysis (200-300 ms).

**Fig. 9.**
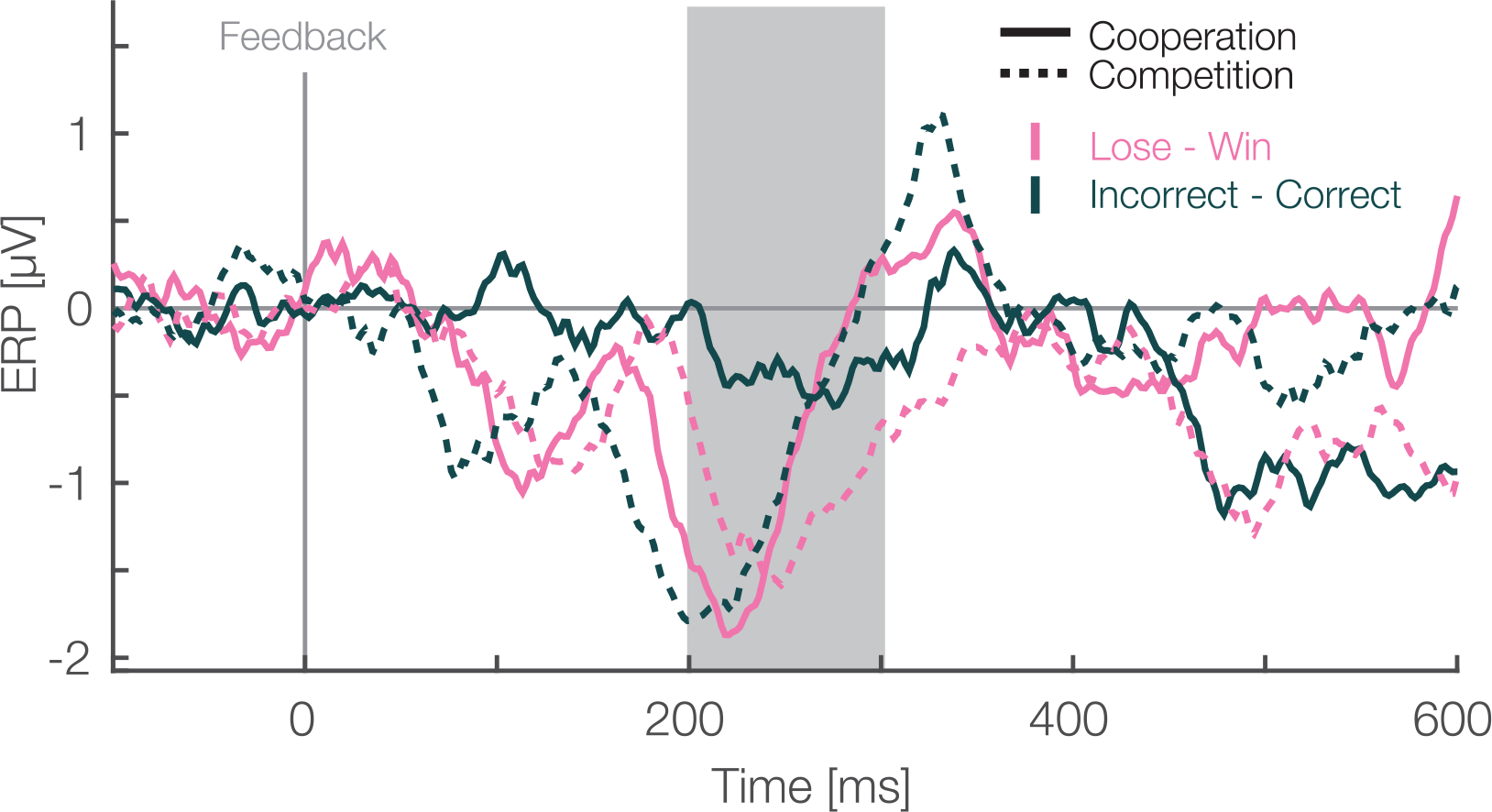
Feedback locked difference waveforms at pooled electrode sites (F1, Fz, F2, FC1, FCz, FC2). Data are average referenced. Pink and green colors represent the monetary outcome, i.e., lose-win and incorrect-correct trials respectively. Solid and dashed lines represent cooperative and competitive situations. The gray box shows the preselected time window used for the confirmatory statistical analysis (200-300 ms).

**Fig. 10.**
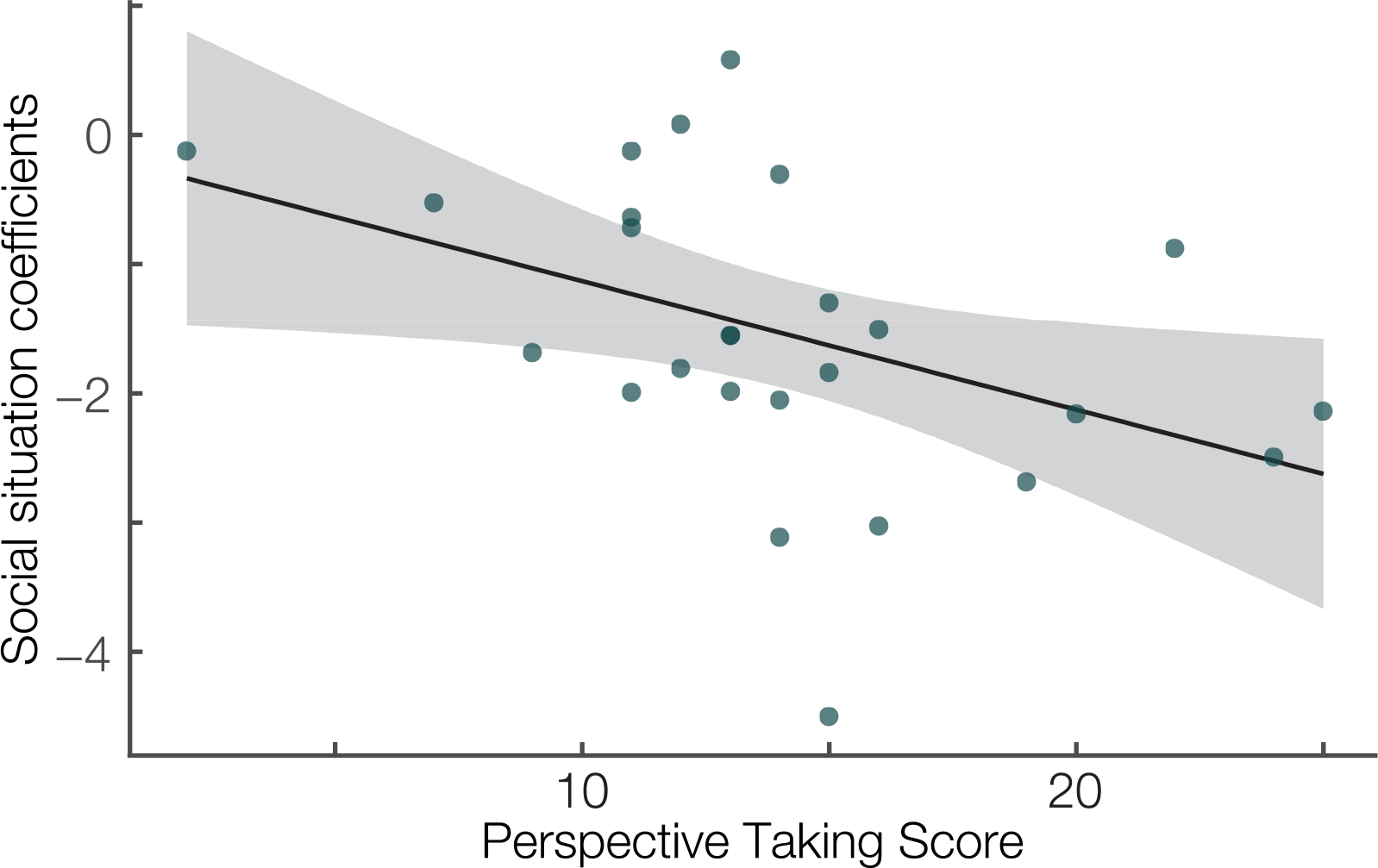
Correlation between the Perspective taking subscale of the Interpersonal Reactivity Index (Davis, 1983) (x-axis) and the single-subject linear mix model estimates of the social situation effect (y-axis). Spearman’s *r* = −.54, *p* = .005. Linear fit with 95% confidence interval.

**Fig. 11.**
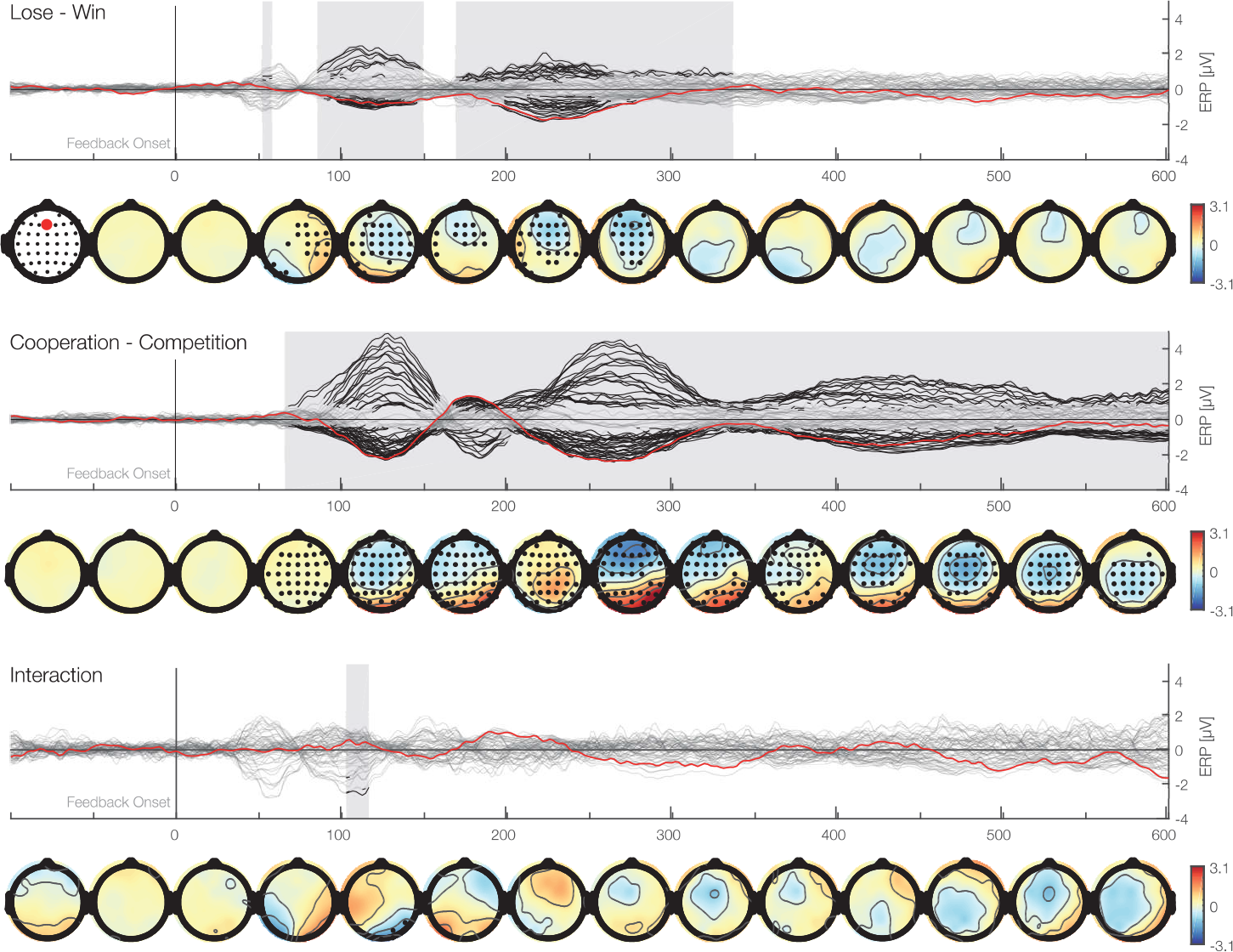
Time-series plots of the EEG amplitudes of the main factors and interaction for each electrode aligned to the feedback stimulus. First row (butterfly plot) shows time against activity of all electrodes. In addition, the Fz electrode is marked in red. Black marked clusters are significant under a TFCE permutation p-value of 0.05. TFCE corrects for multiple comparisons over time and electrodes. Second row shows topographical plots representing the mean amplitudes averaged over 50 ms bins. Black marked electrodes represent significant channels.

**Fig. 12.**
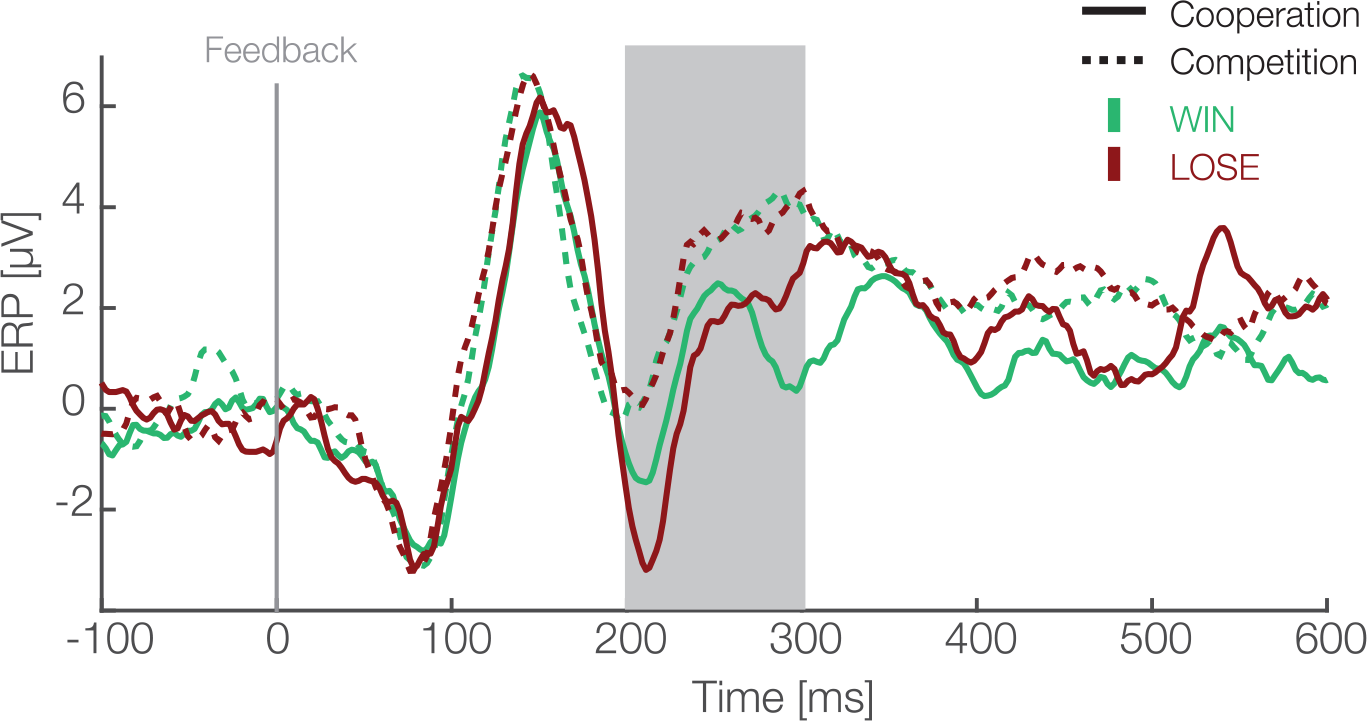
Feedback locked averaged difference waves between experimental and control data for 5 participants pooled at electrode sites (F1, Fz, F2, FC1, FCz, FC2) are shown. In experimental data green and red colors represent the outcome, i.e., win and lose trials respectively, while solid and dashed lines represent cooperative and competitive situations. Difference between social situations and outcomes is visible after subtraction of electrophysiological response to identical visual stimuli without any content. This suggests that our results represent differences in experimental manipulations but not visual properties of stimuli.

## Conflict of Interest Statement

The authors declare that the research was conducted in the absence of any commercial or financial relationships that could be construed as a potential conflict of interest.

## Author Contributions

Study design (ACz, BW, PK), data collection (ACz), data analysis (ACz, BE), draft and revisions of manuscript (ACz,BE,BW,PK).

## Acknowledgements

We gratefully acknowledge the support by the European Commission Horizon H2020-FETPROACT-2014 641321—socSMCs, DFG-funded Research Training Group “Situated Cognition” (GRK 2185/1) and the Deutsche Forschungsgemeinschaft (DFG) Open Access Publishing Fund of Osnabrück University. Moreover, we would like to thank Anna Lisa Gert for her help with preprocessing EEG data as well as following students, who collected the data: Chiara Carrera, Marketa Becevova, Maria Sokotushchenko, Greta Häberle and Susanne Schuberth.

